# A new species of deep-water *Lethrinops* (Cichlidae) from Lake Malawi

**DOI:** 10.1101/2022.04.29.489986

**Authors:** George F. Turner

## Abstract

A new species of cichlid fish, *Lethrinops atrilabris* is described from specimens collected by trawling at a depth of around 90m off Monkey Bay, southern Lake Malawi. It is assigned to the genus *Lethrinops* on the basis of its vertical flank barring, lack of enlarged cephalic lateral line canal pores and the form of the lower jaw dental arcade. It can be distinguished from congeneric species by its male breeding dress of contrasting flank barring and dark ventral surface, most strikingly on the lips, throat and chest, its relatively small known maximum size (<75mm SL), large eyes (38-41% head length), laterally compressed body (depth 2.5-2.7 times max head width) and lower gillraker count (13-14).

## 1. INTRODUCTION

Lake Malawi hosts an enormous number of endemic cichlid fishes, in one recent guide, estimated to be over 800 species (Konings, 2016). Although this extraordinary adaptive radiation is of great interest to evolutionary biologists, conservationists, fishing communities and aquarium fish enthusiasts, the rate of species description is slow and many species – even some well-known ones - remain undescribed, rendering them ineligible to receive IUCN redlisting, or incorporation into standard reference systems such as FishBase, GBIF etc.

The genus *Lethrinops*, as currently understood, comprises 22 described and many undescribed species of sediment-sifting cichlid fishes endemic to the Lake Malawi and Upper/Middle Shire River catchments. They are characterised by the shape of their lower jaw dental arcade, in which the outer row curves in posteriorly to end abruptly behind the inner row(s), if present (Trewavas 1931, Turner 1996, Ngatunga & Snoeks 2004). In the majority of Malawi endemics, the outer tooth row continues in a relatively straight line, often dwindling to a few small widely-spaced teeth: referred to as the ‘Haplochromis’-style, as many of these species were formerly assigned to the genus *Haplochromis*. Two other genera of Lake Malawi cichlids share the ‘*Lethrinops*-type’ dentition and were separated from *Lethrinops* by Eccles & Trewavas (1989) on the basis of shared derived characters: the large long-snouted *Taeniolethrinops* species were reported to have an oblique-striped flank pattern, while the small short-snouted *Tramitichromis* have a distinctive lower pharyngeal bone shape and few widely-spaced gillrakers. *Lethrinops-style* dentition is also shown by *Ctenopharynx pictus*, which is placed in its genus on the basis of traits shared with the other two *Ctenopharynx* species. Therefore, the genus *Lethrinops* is currently defined by a single trait that appears to have evolved repeatedly and by the absence of other presumed derived traits. Perhaps not surprisingly, it has been proposed that it is comprised of two or more groups of species that are not particularly closely-related that can be roughly characterised as ‘shallow-water’ and ‘deep-water’ species, comprised of 12 and 10 described species respectively (Ngatunga & Snoeks 2004).

The aim of the present study is to formally describe an additional deep-water species conforming to the current definition of the genus *Lethrinops* Regan 1922 (by Eccles & Trewavas 1989), known informally as *Lethrinops* ‘black chin’ (Turner, 1996).

## 2. MATERIALS AND METHODS

Specimens were obtained from a research trawl survey carried out by the Monkey Bay Fisheries Research Station (now known as the Fisheries Research Unit, FRU) of the Malawi Government, using the trawler Ethelwynn Trewavas, in 1992, intended to estimate standing stocks of food fishes. The majority of the catch was sold for human consumption, but on this occasion, a few specimens were preserved for research. These were already dead when selected and were pinned and photographed before being preserved in formalin, later being washed and transferred to 70% ethanol for long-term preservation. Counts and measurements were carried out following the methods of Snoeks (2004), using digital callipers and a low power magnifying desk lamp and various eye pieces (loupes).

Comparison with similar species was based on published (re-)descriptions, largely in Eccles & Trewavas (1989) for *Lethrinops* and *Ctenochromis* and Hanssens (2004) for *Placidochromis* along with re-examination of some of the type material, along with specimens held at Bangor University collected from 1990-2017. Direct comparisons of morphometric ratios were avoided as diagnoses resulting from generally small samples of type specimens rarely persist when larger numbers of specimens are examined, particularly when representing a fuller size range. In the author’s experience differentiation of Lake Malawi haplochromines is better achieved by overall appearance, aided by verbal descriptions in combination with meristics, character states such as dentition and male breeding dress.

### Ethical Statement

The study did not involve live animals, as it used preserved specimens that were collected already dead from a trawl catch carried out as part of a Malawi Government research survey.

## 3. RESULTS

### *Lethrinops atrilabris* sp. nov

urn:lsid:zoobank.org:pub:101AB870-D407-416D-85F7-D61D73714064

Holotype: BMNH 2022.4.20.1, male, 72.0mm SL, collected from trawl catch NE of Monkey Bay, at a reported depth of 84-94m, 13^th^ April 1992.

Paratypes: BMNH 2022.4.20.2-7, six males 66.2-72.9mm SL, collected with holotype.

Diagnosis: the lower jaw dentition ‘*Lethrinops*-type’. Mature males with a melanic pattern of strongly contrasting dark vertical flank bars on a pale background, and a dark area on the jaws and the underside of the head and chest. In addition, the species can be identified by its relatively small adult body side (not known to exceed 73mm SL), large eye, short, rounded snout, ventrally-placed mouth, 13-14 ceratobranchial gill rakers and laterally compressed body.

Comparisons: The male’s melanic pattern of strongly contrasting vertical flank bars is not exhibited by any known species of *Ctenochromis, Taeniolethrinops* or *Tramitichromis*. Among the described *Lethrinops* species, males of the shallow-water group (sensu Ngatunga & Snoeks 2004) do not show such strong vertical flank barring and tend to be less deep-bodied and laterally compressed and confined to shallower water (generally <50m, compared to 84-94m for *L. atrilabris*). This group comprises *Lethrinops albus* Regan 1922, *Lethrinops auritus* (Regan 1922), *Lethrinops furcifer* Trewavas 1931, *Lethrinops lethrinus* (Günther 1893), *Lethrinops leptodon* Regan 1922, *Lethrinops lunaris* Trewavas 1931, *Lethrinops macrochir* (Regan 1922), *Lethrinops macrophthalmus* (Boulenger 1908), *Lethrinops marginatus* Ahl 1927, *Lethrinops microstoma* Trewavas 1931, *Lethrinops parvidens* Trewavas 1931, *Lethrinops turneri* Ngatunga & Snoeks 2003 and a number of undescribed species. Among the remaining, ‘deep-water’ *Lethrinops* species are 10 described species. *Lethrinops atrilabris* has a greater number of lower gillrakers (13-14) than *Lethrinops christyi* Trewavas 1931 (8-9), *Lethrinops longipinnis* Eccles & Lewis 1978 (9-10) and *Lethrinops altus* Trewavas 1931 (10-11). These three species can further be distinguished by their head and jaw shape: *L. christyi* has small pointed jaws and concave upper profile of snout v larger jaws set low on a rounded head profile in *L. atrilabris; L. longipinnis* has a much longer snout; *L. altus* has hooked maxillae, showing a markedly curved lower profile, in contrast to the straight maxillae in *L. atrilabris*. *Lethrinops atrilabris* has fewer lower gillrakers (13-14) than *Lethrinops micrentodon* (Regan 1922) (15-19), *Lethrinops gossei* Burgess & Axelrod 1973 (18-19), *Lethrinops stridei* Eccles & Lewis 1977 (19-23), *Lethrinops macracanthus* Trewavas 1931 (21-24) and *Lethrinops microdon* Eccles & Lewis 1977 (24-29). *Lethrinops mylodon* Eccles & Lewis 1979 generally has fewer lower gillrakers (10-14 v 13-14 in *L. atrilabris*) and also differs in having a very heavily-built lower pharyngeal bone with stout molariform teeth (v lightly-built, with small slender teeth in *L. atrilabris*) and in attaining a much larger size (>200mm SL v <80 mm SL in *L. atrilabris). Lethrinops longimanus* Trewavas 1931 generally has a higher count of lower gillrakers:15-19 according to Eccles & Lewis 1979, although Eccles & Trewavas (1989) give 14 as the lower limit, v 13-14 in *L. atrilabris. Lethrinops longimanus* can also be distinguished by its larger maximum size (150mm SL v <80mm SL) and male breeding dress of a bronze colour, weakly barred v the strongly barred black and silver of *L. atrilabris*.

The dental arcade trait can be difficult to see without a powerful microscope and appropriate lighting, so this trait is of little use to fieldworkers. Other deep-bodied, deep-water species with similar barred patterns are presently classed in the genera *Alticorpus, Aulonocara* and *Placidochromis*. Members of the first two genera are distinguished by having much larger cephalic lateral line pores, particularly on the underside of the head, that other Malawian cichlids, including *Lethrinops*. Distinguishing *Placidochromis* species can be more problematic, as these lack this diagnostic trait. A number of deep-water species were described by Hanssens in 2004, several superficially resembling *L. atrilabris*. From these, *L. atrilabris* can be distinguished by its lower-arch gillraker counts (13-14), which are lower than those of *Placidochromis chilolae* Hanssens 2004 (14-16), *Placidochromis lukomae* Hanssens 2004 (14-18), *Placidochromis nigribarbis* Hanssens 2004 (16-18), *Placidochromis obscurus* Hanssens 2004 (18-21) and higher than *Placidochromis domirae* Hanssens 2004 (8-9), *Placidochromis koningsi* Hanssens 2004 (10), *Placidochromis msakae* Hanssens 2004 (12), *Placidochromis pallidus* Hanssens 2004 (11-12), *Placidochromis rotundifrons* Hanssens 2004 (11) and *Placidochromis turneri* Hanssens 2004 (9-10). Other species in the genus can be differentiated quite readily on physical appearance, such as having a shallower body, smaller eyes, a longer, more pointed snout, larger jaws or a mouth in a more terminal position or more upwardly-angled (see illustrations in Hanssens 2004 or Konings 2016).

#### Description

Body measurements and counts in table 1. *Lethrinops atrilabris* is a small (<80mm SL) laterally-compressed (maximum body depth 2.5-2.7 times maximum head width) cichlid fish with a short, rounded snout (27-32% HL), small mouth low down on the head and very large eyes (38-41% HL). To date, only mature males have been identified and these have conspicuously barred flanks and a black underside to the head and chest (Figure 1).

**Figure 1:**
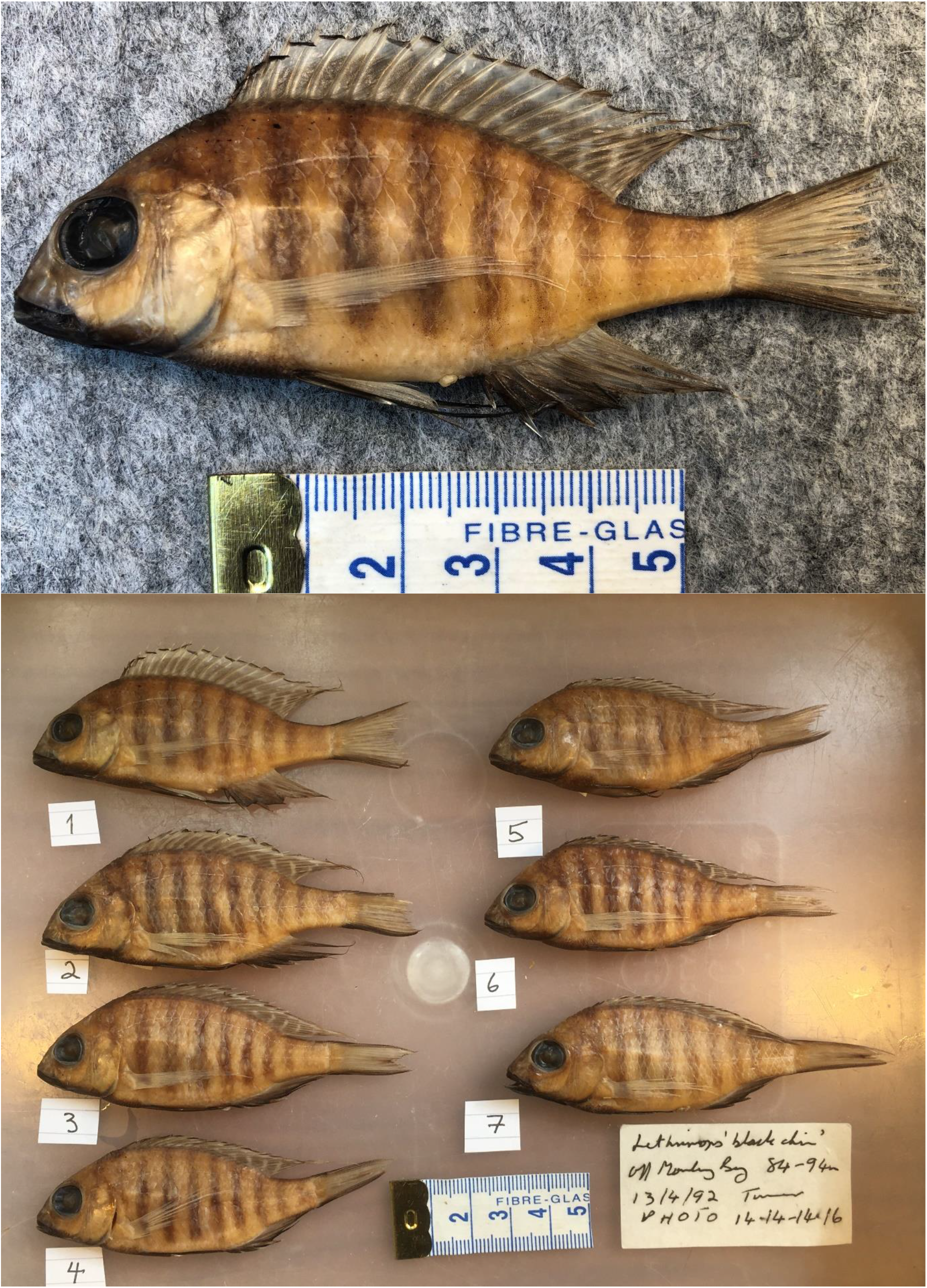
*Lethrinops atrilabris* sp. nov. Above: holotype: BMNH 2022.4.20.1, male, 72mm SL, collected from trawl catch NE of Monkey Bay, at a reported depth of 84-94m, 13^th^ April 1992; Below: the full type series, holotype labelled 1, collecting information as holotype.

**TABLE 1.** Morphometric and meristic characters of *Lethrinops atrilabris*.

|  | Holotype | Paratypes (n=6)<br>mean & range |
| --- | --- | --- |
| Standard Length (mm) | 72.0 | 69.2 (66.2-72.9) |
| **As % SL** |  |  |
| Maximum Body Depth | 38.6 | 39.2 (38.1-39.8) |
| Head Length | 32.9 | 32.9 (32.1-33.6) |
| Dorsal-Fin Base Length | 57.2 | 57.0 (53.7-58.8) |
| Anal-Fin Base Length | 18.8 | 17.4 (16.7-18.4) |
| Predorsal Length | 39.6 | 37.5 (36.6-38.1) |
| Preanal Length | 64.4 | 66.5 (65.4-69.2) |
| Prepectoral Length | 32.8 | 33.9 (32.3-34.8) |
| Prepelvic Length | 39.9 | 39.8 (38.1-41.5) |
| Caudal-Peduncle Length | 16.7 | 16.2 (15.6-16.9) |
| Caudal-Peduncle Depth | 12.1 | 11.9 (11.6-12.2) |
| **As % Head Length** |  |  |
| Head Width | 44.7 | 46.1 (45.0-47.5) |
| Interorbital Width | 22.8 | 23.9 (22.1-27.4) |
| Snout Length | 32.1 | 29.1 (26.7-30.4) |
| Lower Jaw Length | 39.2 | 39.0 (37.2-41.1) |
| Premaxillary Pedicel Length | 27.0 | 25.3 (24.2-26.1) |
| Cheek Depth | 16.9 | 17.3 (16.6-18.2) |
| Eye Diameter | 40.9 | 39.8 (38.3-40.8) |
| Lachrymal Depth | 21.1 | 21.1 (20.4-22.9) |
| **Ratios** |  |  |
| Body Depth/Head Width | 2.62 | 2.58 (2.51-2.67) |
| Caudal-Peduncle Length/Depth | 1.38 | 1.36 (1.30-1.43) |
| **Counts** |  |  |
|  | Holotype | Paratypes<br>range |
| Upper Gillrakers | 5 | 4-5 |
| Lower Gillrakers | 14 | 13-14 |
| Dorsal Fin Rays | XVI, 9 | XV-XVI, 9-10 |
| Anal Fin Rays | III, 7 | III, 7-9 |
| Longitudinal Line Scales | 32 | 31-34 |
| Cheek Scales | 2 | 2 |

The size range of the seven specimens is 66-73mm SL. As all specimens collected showed clear evidence of male breeding dress, it can be assumed that all are adult males, probably collected on a breeding ground. In haplochromine cichlids, the largest males are typically larger than the largest females, and there is not usually a great deal of variation in the size of adult males on breeding grounds. As the specimens were collected from an unselective trawl catch along with many much larger individuals of other species, it seems likely that the maximum adult size of this species is less than 80mm SL, at least in the SE Arm of the lake.

All specimens relatively deep-bodied, laterally compressed, deepest part of body generally well behind first dorsal fin spine. Anterior upper lateral profile convex and gently curving, without a sharp inflection in curve above the eye. Lower anterior lateral profile also gently curving, so that tip of snout lies well above insertion of pelvic fins. Mouth relatively small, low on head, slightly upwardly-angled, snout well below horizontal plane from bottom of eye. Eye extremely large, circular, generally appearing more or less touching anterior upper lateral head profile. Lachrymal much wider than deep, 5 openings.

Flank scales weakly ctenoid, cteni becoming reduced dorsally, particularly anteriorly above upper lateral line, where they transition into a cycloid state. Scales on chest are relatively large, gradually transitioning in size from larger flank scales, as is typical in non-mbuna Malawian endemic haplochromines (Eccles & Trewavas 1989). A few small scales scattered on the proximal part of caudal fin.

Cephalic lateral line pores inconspicuous, flank lateral line shows the usual cichlid pattern of separate upper and lower portions.

Pectoral fin very long when intact, extending well past first anal spine. Pelvic fins extend past vent in all specimens and past first anal spine in some: this may be a sexually dimorphic trait, with female haplochromines often having shorter pelvic fins. Tips of dorsal and anal fins also prolonged, extending well past the plane through base of caudal fin in some specimens- again probably a sexually dimorphic trait, exaggerated in males. Tailfin crescentic.

Lower jaw relatively small, with thin mandibular bones, but not flattened as it is in some *Placidochromis*, such as *P. hennydaviesae*. Jaw teeth small, short and erect. Outer series in both upper and lower jaw largely unequally bicuspid, becoming more equally bicuspid posteriorly, notably in upper jaw. A single inner series of very small tricuspid teeth.

Lower pharyngeal bone small, lightly-built, Y-shaped, and carries small, short, laterally compressed slightly hooked, blunt, simple teeth. Middle-lying 5-6 teeth on each side of posterior row slightly larger than others, but molarization lacking. About 12 teeth in midline row and about 20 on each side on posterior row. Gill rakers simple, erect, fairly long and well-spaced, with few, if any, reduced to small stubs near anterior part of arch.

Colouration of females and immatures is unknown, but from experience of other species from this habitat, can be expected to be countershaded, sandy-coloured dorsally, with silvery flanks and probably faint vertical flank bars. All known specimens appear to be males in breeding dress. Colour notes based on a photograph of a freshly collected type specimen and an additional specimen collected in 2016, but not yet located in the collection at Cambridge University (Figure 2). Strong dark brownish vertical flank bars on silvery-white background: 6 bars under dorsal fin, 2 more on caudal peduncle and 1-2 on nape. Head dark brown on upper surface, but paler laterally, sometimes with a dark lachrymal mark running from eye toward the mouth. Eye golden brown, darker along the axis of lachrymal stripe. Lips, lower jaw, throat and chest are black. Dorsal fin dark golden-brown, with series of irregular white spots or oblique stripes angled forwards from base, with broad black margin and broader white submarginal band. Pectoral fins translucent, but brownish-tinted. Pelvic fins black, fading to dark grey on posterior rays. Anal fin black, fading to dark grey basally and marked with irregular yellowish spots and stripes. Caudal fin with dark grey to black upper and lower margins, but otherwise dark golden-brown with three thin irregular vertical white bands.

**Figure 2:**
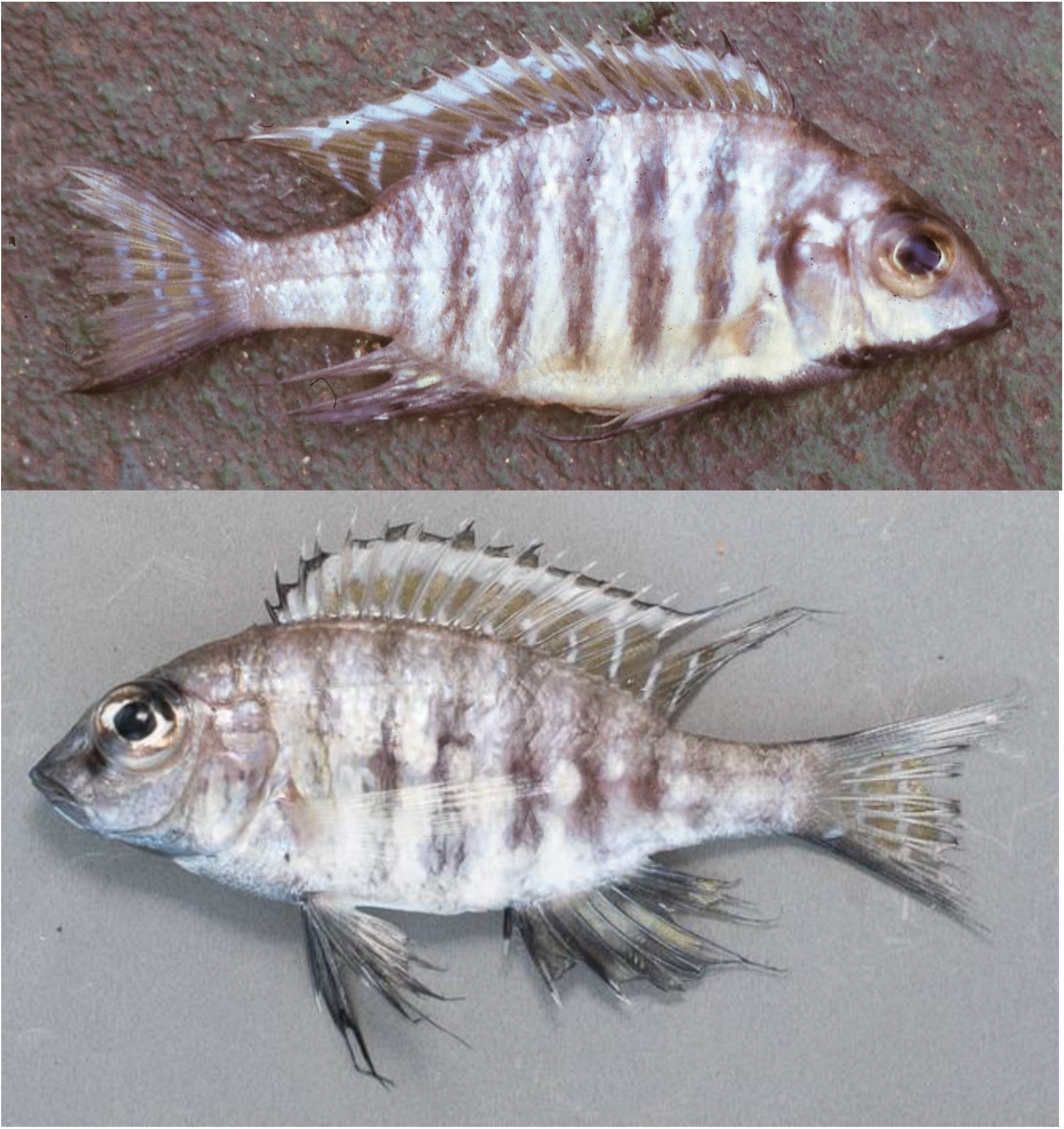
*Lethrinops atrilabris* sp. nov. Fresh coloration. Above: one of the type specimens photographed shortly after capture. Below: probable *L. atrilabris*, collected from trawl catch at 95-105m depth, East of Domwe Island, SE Arm, 4th March 2016. Cambridge University collection, identification not confirmed.

#### Distribution

Positively known only from the type locality, in the SE Arm of Lake Malawi, NE of Monkey Bay, at a reported depth of 84-94m. A photograph of a possible specimen of this species was taken from a trawl catch at 95-105m East of Domwe Island, SE Arm, 4th March 2016. These two localities are close together, as are the depths.

#### Etymology

‘Atri-’ from plural of the adjective ‘ater’ (Latin) = black + ‘labris’ from plural of labrum (Latin)= lip, in reference to the black lips of the males in breeding dress.

## 4. DISCUSSION

The cichlid genus *Lethrinops* is endemic to Lake Malawi and its catchment and the outflowing Shire River, its expansion in Lake Malombe and continuation to the biogeographic barrier represented by the falls on the middle Shire, notably the Kapichira rapids, below which the fish fauna is essentially lower Zambezian (Tweddle & Willoughby 1979). Originally defined by Regan (1922) based on its dentition-principally in having small, weak teeth in narrow bands-the genus originally included just 4 species, including the type *L. lethrinus*. Trewavas (1931) revised the genus, her definition emphasising the semicircular shape of the lower jaw dental arcade, and increasing the number of included species to 23. The revision by Eccles & Trewavas (1989) split the genus into three. Five small, short-snouted species were moved into *Tramitichromis*, characterised by the shape of the lower pharyngeal bone, in which the upper margin of the blade is turned sharply downwards and the anterior end of the pharyngeal dental arcade is broad and rounded. In addition, four large, long-snouted species were grouped into *Taeniolethrinops*, characterised by having an oblique dark stripe on the flanks of females and immature fishes (although not all species actually seem to show this in my experience). Thus, *Lethrinops* was left without any defining synapomorphy: characterised by its dental arcade-shared with *Tramitichromis* and *Taeniolethrinops-* but lacking the diagnostic traits of the latter two genera.

Early molecular studies using mitochondrial DNA restriction fragment analyses placed the deep-water *Lethrinops gossei* in a surprising grouping with the mbuna species, along with a number of *Aulonocara* species, and not with the major ‘Haplochromis’ or ‘sand-dweller’ group from sandy or muddy habitats (Moran et al. 1994). However, later studies placed a number of shallow-water *Lethrinops* and a *Taeniolethrinops* species in the ‘sand-dweller’ group, suggesting the genus to be polyphyletic (Joyce et al. 2011, Genner & Turner 2012). In addition, the deep-water species were shown to have an affinity with *Alticorpus* and some deep-water *Placidochromis* species. Early nuclear gene analyses presented rather inconsistent pictures, but whole genome sequencing (Malinsky et al. 2018; Masonick et al. 2022) has continued to support the distinctness of the deep-water and shallow-water *Lethrinops* species, and the affinity of the former to *Aulonocara* and *Alticorpus* (deep-water *Placidochromis* were not investigated).

On the basis of the emerging mitochondrial data, Ngatunga and Snoeks (2004) informally split the genus into deep-water and shallow-water groups, with the type species, *Lethrinops lethrinus* clearly a member of the latter, suggesting that the deep-water species will be in need of a new generic classification. However, this has yet to be attempted and at present the distinction is unclear.

Generally, the deep-water species mostly occur at depths of 50m or more and seem to be relatively deep-bodied and laterally compressed. Males in breeding dress tend to express strong vertical barring on their flanks, as do species of *Alticorpus*, *Aulonocara* and *Placidochromis* from the same habitat, while shallow-water *Lethrinops* males are usually unbarred or weakly-barred with a range of bright colours including red, orange, yellow, blue and green: see illustrations in Konings (2016), for example. A few species, such as *L. altus, L. christyi, L. longimanus, L. longipinnis* and *L. micrentodon* are more problematic, with forms exhibiting a mix of traits, and often being found at depths of 20-60m. However, *Lethrinops atrilabris* is unambiguously a member of the deep-water group, with its strongly barred males and relatively deep, laterally compressed body. The species shows superficial similarities to a number of species of the genus *Placidochromis*, which also includes a number of deep-water, vertically-barred species. From these, it can be distinguished by the shape of the lower jaw dental arcade (Hanssens 2004). However, it is not clear whether this trait really has much phylogenetic significance: this will probably require extensive whole genome sequencing and phylogenetic analysis.

An additional case of evolution of the *Lethrinops*-style dentition appears to have occurred in *Ctenopharynx pictus*, which, like known species of *Lethrinops, Taeniolethrinops* and *Tramitichromis*, is a sediment-sifting species. Eccles & Trewavas (1989) placed this species in *Ctenopharyx* on the basis of its spotted melanin pattern, large number of gillrakers and ‘weak’ jaws and dentition. This classification is supported by recent genome-wide analysis (Masonick et al. 2022), although the specimen is mistakenly labelled as *‘Otopharynx pictus’*, possibly following a period of usage of *Ctenopharynx* as a subgenus of *Otopharynx* in some publications in the 1990s (e.g. Konings 1990).

## ACKNOWLEDGEMENTS

I am grateful to all my colleagues and collaborators in Malawi during the late 1980s and early 1990s who helped me with collecting and identifying the fishes we sampled during the ODA (now DFID)-funded Lidole Project and the FAO Chambo Fisheries Project, carried out in collaboration with the Malawi Government Fisheries Research Unit at Monkey Bay, and I thank James Maclaine at the Natural History Museum in London for curating the specimens.

